# Low extent of sex-based venom variation in Earth’s most widespread viper, the common adder (*Vipera berus*)

**DOI:** 10.1101/2025.03.27.645687

**Authors:** Lennart Schulte, Miguel Engelhardt, Johanna Eichberg, Alfredo Cabrera-Orefice, Harry Wölfel, Maik Damm, Ignazio Avella, Benno Kreuels, Kornelia Hardes, Johannes A. Eble, Andreas Vilcinskas, Tim Lüddecke

## Abstract

Snake venom is an ecologically critical functional trait, primarily applied for foraging and accordingly shaped by selective pressures. Recent insights underpinned the high variability of snake venoms down to the intraspecific level, with regional, ontogenetic, and seasonal variation being mostly investigated. In contrast, sex-based venom variation has received considerably less attention so far, and its influence on venom compositions in vipers is virtually unknown. The common adder (*Vipera berus*) is a promising model species to explore this subject because of a described sexual dimorphism and its wide distribution, which promises a noteworthy degree of adaptability and venom plasticity. Here, we tested for sex-based venom variation in Central European *V. berus* by comparing venom profiles and bioactivity. Proteomics, paired with SDS-PAGE and RP-HPLC, revealed highly similar venom profiles. Likewise, phospholipases A_2_ and proteases bioactivity profiling, and bioassays targeting the coagulation cascade and mammalian cell lines revealed similar activity spectra. Hence, our analysis does not show a noteworthy extent of sex-based intraspecific venom variation in *V. berus*. We further discuss our data in light of the species’ venom profile at larger geographic scales, its clinical relevance, and the need for more standardized approaches when investigating venom variability. Our work provides novel insights into the venom biology of Earth’s most widespread viper, and serves as a foundation upon which future works can build.

## 2 Introduction

Snake venoms are complex mixtures of several bioactive molecules, mainly proteins and peptides, able to alter the physiological balance of the organism into which they are injected (Casewell *et al*., 2013). Extensive evidence suggests that the main selective driver of snake venom evolution and variation is predation (Daltry *et al*., 1996b; Barlow *et al*., 2009; Holding *et al*., 2021). Furthermore, various factors have been shown to affect the type and abundance of snake venom compounds, thus determining remarkable compositional and functional variation at all taxonomic levels. Indeed, while interspecific venom variation has long been acknowledged, intraspecific variation has recently started to be recognized and investigated (Casewell *et al*., 2020). Among the drivers of intraspecific venom variation identified so far are differences in geographic origin (Zancolli *et al*., 2019; Avella *et al*., 2023), life history stages (Cipriani *et al*., 2017; Avella *et al*., 2022; Ferreira-Rodrigues *et al*., 2024), and sex of the specimens considered (Menezes *et al*., 2006; Ferreira-Rodrigues *et al*., 2024). Sex-related venom variation is expected in taxa that employ their venom for intraspecific competition and/or displaying sexually dimorphic feeding or dispersal ecologies (Schendel *et al*., 2019; Lüddecke *et al*., 2022). Considering snakes, evidence of venom being used for intraspecific competition is currently lacking, and male and female conspecifics typically present virtually identical diets (Madsen and Shine, 1994; Bonnet *et al*., 1998; Avolio *et al*., 2006; Do *et al*., 2023). Therefore, while intraspecific venom variation can often be explained by differences in diet and/or prey availability (e.g., between snakes from different localities or age), the occurrence of sex-related venom variation remains a conundrum.

To date, a limited number of studies have investigated this question within members of the family Viperidae, with pit vipers (subfamily Crotalinae) being the major model group. For instance, Menezes *et al*. (2006) detected variation in venom composition and activities between male and female *Bothrops jararaca* siblings, suggesting it to be genetically inherited and imposed by evolutionary forces. Similarly, Zelanis *et al*. (2016) demonstrated the presence of sex-related compositional and functional variation in the same species. Furthermore, Furtado *et al*. (2006) proposed that the observed differences in composition and bioactivity between females and males might be attributed to sexual dimorphism associated with different diets. Studies in *Bothrops moojeni* also indicate a pronounced sex-based venom variation within this species (Hatakeyama *et al*., 2021; Ferreira-Rodrigues *et al*., 2024), alongside documented differences in diet where males and juveniles feed mainly on ectotherms, while the considerably larger adult females prefer mammalian prey (Nogueira *et al*., 2003). Similar claims have been made across a number of other members of the genus *Bothrops* (Machado Braga *et al*., 2020; Hatakeyama *et al*., 2020; Kallel *et al*., 2024). However, studies in *Bothrops jararacussu* (Aguiar *et al*., 2020) and *Bothrops asper* (Gómez *et al*., 2021) found no significant differences between male and female venoms, despite marked sexual dimorphism.

Furthermore, proteomics on the venom of *Tropidolaemus wagleri*, a species of pit viper presenting remarkable sexual size dimorphism, did not reveal noteworthy differences in venom composition and lethality between sexes (Tan *et al*., 2017). Besides pit vipers, venom variation has been sporadically investigated in other viperid snakes. For instance, several SDS-PAGE profiling studies on Old World vipers (subfamily Viperinae) showed at least subtle differences between male and female venoms (Marsh and Glatston, 1974; Mebs and Kornalik, 1984). Recent proteomic studies on the genus *Vipera* (Petras *et al*., 2019; Avella *et al*., 2023; Lakušić *et al*., 2025a) have identified varying degrees of sex-based differences. However, these differences were generally considered to have an insignificant impact on venom composition, although bioactivity assessments remain lacking. Given this, it is clear that the underlying drivers and the extent of these differences require further investigation.

The common adder (*Vipera berus*) represents a promising model species to investigate sex-based snake venom variation. It is a medically important snake with the widest distribution in the world, ranging from the United Kingdom to North Korea and southeastern Russia (Figure 1, A), where it occurs in a variety of habitats (Speybroeck *et al*., 2016, 2020; Geniez, 2018; World Health Organisation, 2024). As in several other vipers, *V. berus* presents a marked ontogenetic shift in diet, with juveniles mostly feeding on ectotherm prey (e.g., reptiles and amphibians), and gradually incorporating more endotherms (e.g., small mammals and birds) in their diet as they grow (Otte *et al*., 2020; Samsonov *et al*., 2022). Furthermore, *V. berus* exhibits sexual dimorphism in color and size (Figure 1, B), with females being generally longer and bulkier than males, likely to facilitate reproduction (Forsman, 1991; Madsen and Shine, 1994). To date, whether this sexual dimorphism is associated with differences in diet composition between male and female common adders remains inconclusive (Forsman, 1991).

**Figure 1.**
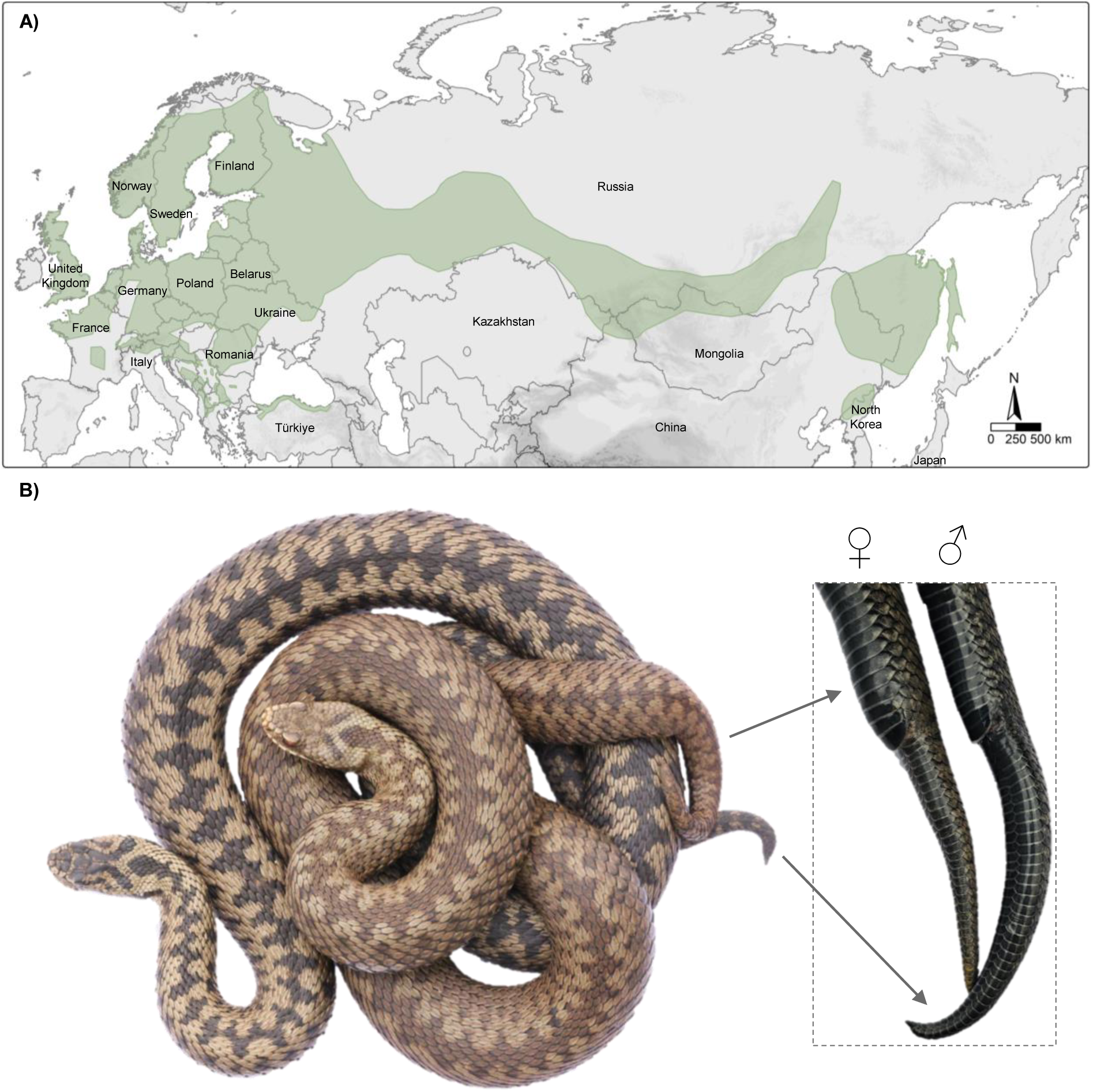
The common adder, *Vipera berus*. Geographic distribution of *V. berus* (green), based on data from the International Union for Conservation of Nature (IUCN; accessed 20 March 2024). *(B)* Representative male (left) and female (right) *V. berus* specimens, illustrating sexual dimorphism in coloration and tail morphology. The male exhibits a whitish body with a distinct dark zigzag dorsal pattern, whereas the female displays a pale brownish body with a less contrasting brownish zigzag pattern. Tail morphology differences are highlighted. The distribution was plotted using R version 4.2.2 (R Core Team, 2024). Credits: Harry Wölfel

Due to its wide distribution, *V. berus* is frequently involved in snakebite incidents and is thus listed among the snakes of the highest medical relevance in Europe (Hermansen *et al*., 2019; Di Nicola *et al*., 2021). Proteomic studies identified phospholipases A_2_ (PLA_2_s), snake venom metalloproteinases (svMPs), and snake venom serine proteases (svSPs) as major enzymatic components within the venom of this species (Ramazanova *et al*., 2008; Latinović *et al*., 2016; Bocian *et al*., 2016; Al-Shekhadat *et al*., 2019; Damm *et al*., 2021). Intriguingly, several studies have highlighted the presence of considerable variation in the diversity and abundance of venom compounds within *V. berus*, indicating differences related to several factors, such as regional, ontogenetic, and individual variation (Mebs and Langelüddeke, 1992; Nedospasov and Rodina, 1992; Malina *et al*., 2017). Similar patterns were observed in venoms of *V. berus* subspecies, such as *V. berus barani* (Damm *et al*., 2024) and *V. berus nikolskii* (Kovalchuk *et al*., 2016), as well as closely related species e.g. *Vipera seoanei* (Avella *et al*., 2023), *Vipera kaznakovi* (Petras *et al*., 2019), and *Vipera ammodytes* (Lakušić *et al*., 2025a; b). Taken together, this suggests that the extent of intraspecific venom variation within *V. berus* is likely underestimated, and the underlying factors are yet to be linked to conclusive patterns. This particularly concerns sex-based venom variation, which may occur within this species as per its sexual dimorphism.

Here, we present the first in-depth investigation of sex-based variation in *V. berus* venom. We tested for qualitative differences between the pooled venoms of male and female adult specimens of the same region by comparing SDS-PAGE, RP-HPLC profiles, and shotgun proteomes. Furthermore, we screened the bioactivity of the venoms based on previously described dominant enzymatic toxin families (Latinović *et al*., 2016; Bocian *et al*., 2016; Al-Shekhadat *et al*., 2019; Damm *et al*., 2024; Nicolaysen *et al*., 2024), their modes of action, and clinical reports of *V. berus* envenomation (Persson, 2014; Hermansen *et al*., 2019; Di Nicola *et al*., 2021; Siigur and Siigur, 2022). Therefore, we performed bioassays aiming to elucidate protease and PLA_2_ activities of the venoms, as well as the role they play in activation of Factor Xa (FXa)-like, thrombin-like, and plasmin-like blood-coagulation-related enzymes. Finally, we assessed the venoms impact on the cell viability of two different mammalian cell lines (i.e., MDCK II and Calu-3), their effect on the coagulation time of human plasma, and their hemolytic effect on equine erythrocytes.

## 3 Materials and Methods

### 3.1 Venom

Venoms were gifted by members of the “Terrarienclub Bayreuth und Umgebung e.V.”, and were sourced from captive-bred, adult *V. berus* specimens originating from northern Bavaria (Germany). The venoms were non-invasively collected by letting each viper bite a parafilm-covered microtube without applying any pressure to the venom glands. The obtained venom samples were collected individually and stored on dry ice until lyophilization. Individual venoms of six male and six female vipers were weighed and then redissolved in double-distilled water (ddH_2_O) prior pooling by sex. Aliquots were lyophilized and stored at –20 °C until analysis. Specimens’ sex, body size, and dry venom amount are reported in Supplementary Table 1.

### 3.2 Compositional profiling of the venoms

#### 3.2.1 Reducing- and non-reducing SDS-PAGE

Venom profiling based on sodium dodecyl sulfate-polyacrylamide gel electrophoresis (SDS-PAGE) was carried out following the procedure previously applied by our group (see Schulte *et al*., 2023). Therefore, reducing and non-reducing SDS-PAGE with 5 µg of venom was carried out for both venom pools. Raw gel image is provided in Supplementary Figure 1.

#### 3.2.2 RP-HPLC profiling

For reverse-phase high-performance liquid chromatography (RP-HPLC), the protocol applied by Schulte *et al*., 2023 was adjusted as follows: 200 µg of lyophilized venom was reconstituted in ddH_2_O with 5% (v/v) acetonitrile (MeCN) and 1% (v/v) formic acid (HFo) to a final concentration of 10 mg/ml. After centrifugation for 10 min at 12,000 × *g*, the supernatant (20 µL) was measured on a reverse-phase Discovery BIO wide Pore C18-3 column (4.6 × 150 mm, 3 µm particle size; Supelco) operated by an HPLC Agilent 1100 (Agilent Technologies) chromatography system. The column was heated to 40 °C and the following gradient with solvent A (ddH_2_O with 0.1% (v/v) HFo) and solvent B (MeCN with 0.1% (v/v) HFo) at a flow rate of 1 ml/min was used, given at min (B%): 0-5 (5% const.), 5-65 (5 to 45%), 65-75 (40 to 70%), 75-80 (95% const.), and 10 min re-equilibration at 5% B. The chromatograms were detected by a diode array detector (DAD) at λ = 214 nm wavelength (360 nm reference). A previous blank run, injected with 20 µl of ddH_2_O with 5% (v/v) MeCN and 1% (v/v) HFo, centrifuged for 10 min at 12,000 × *g*, was measured under identical parameters and subtracted from the venom profiles. Raw measurements at λ = 214 nm are provided in Supplementary Table 2.

#### 3.2.3 Shotgun proteomics

In preparation for the shotgun mass spectrometry (MS) approach, 50 µg of lyophilized venoms were dissolved to a final concentration of 1.7 µg/µl in an aqueous solution of 6 M Guanidinium hydrochloride (GdnHCl) and 100 mM Tris(hydroxymethyl)aminomethane hydrochloride (Tris/HCl, pH 8.5) for complete denaturation. Further, the venoms were transferred to Protein LoBind tubes (0030108116, Eppendorf), incubated for 30 min at 37 °C with 10 mM Dithiothreitol (DTT) for disulfide reduction, and followed by incubation for 30 min at 22 °C in the dark with 40 mM chloroacetamide for alkylation of free thiols. Afterwards, Trypsin/LysC mix (V5071, Promega) was added at a 1:50 protease-to-protein ratio and incubated for 1 hour at 37°C and 500 rounds per minute (RPM) shaking. The samples were then diluted 1:7 with 50 mM Tris/HCl (pH 8.0) to decrease the GdnHCl concentration below 1 M and to reactivate trypsin. The digestion was continued overnight at 37 °C and 500 RPM shaking and stopped the next day by adding 1.5 % trifluoroacetic acid. The resulting peptides were desalted and cleaned-up using C18-Chromabond columns (730011, Macherey-Nagel). The eluted peptides were dried using a vacuum-concentrator plus (Eppendorf) and redissolved in 20 μl of an aqueous solution with 5% (v/v) MeCN and 0.15% (v/v) HFo, by vortexing. The peptides were transferred to 96-well PCR plates (PCR-96-FS-C, Axygen), sonicated for 5 min in a water bath, and loaded for LC-MS/MS analysis.

Prior to MS, a coupled chromatographic separation of the peptides was performed on an UltiMate 3000 RSLCnano device (Thermo Fisher Scientific). From the prepared venoms, we loaded 5 μl into a PepMap Neo Trap column (Thermo Fisher Scientific) for concentration and desalting, followed by separation using a 50 cm µPAC column (PharmaFluidics, Thermo Fisher Scientific) connected with a TriVersa NanoMate (Advion) robot for chip-based nano electrospray ionization. Throughout the analysis, the analytical column was kept in an oven at 35°C. Peptide elution was performed using a linear gradient of buffer A (HPLC-grade H_2_O with 0.1 % (v/v) HFo) and buffer B (MeCN with 0.1% (v/v) HFo) at flow rate of 0.7 µl/min, given at min (%B): 0-90 (5–35%), 90-100 (35-85%), 100-108 (cons. 85%), and equilibrated for 10 min at 5% buffer B. MS analysis of the peptides was carried out on an Orbitrap Eclipse Tribrid MS (Thermo Fisher Scientific) in positive mode. The NanoMate-assisted electrospray was established by applying 1.7 kV; the source temperature was set to 250 °C. MS2 spectra were obtained in data-dependent acquisition (DDA) with CID fragmentation and data-independent acquisition (DIA) with HCD fragmentation.

We used Xcalibur v4.3.73.11. (Thermo Fisher Scientific) for data acquisition and analysis. Protein identification was performed with PEAKS 12 (Bioinformatics Solution Inc.) and searched against the UniProt database (taxonomy: “serpentes”; status: “reviewed”; “canonical and isoforms”; accessed 11.02.2025). Analyzing the DDA data, precursor ion mass tolerance set to 15 ppm with 3 missed cleavages allowed and carbamidomethylation set as fixed modification. Fragment ion mass tolerance in linear ion trap MS2 detection was set to 0.5 Da, and the false discovery rate was limited to 0.1. For the qualitative analysis, we only considered proteins that were identified with a –10lgP score of at least 15 and at least two unique peptides. For the analysis of the DIA data, the combined spectral library from male and female DDA analysis was used, applying the same database and settings as for the DDA analysis. The detailed list of parameters for data acquisition, analysis, and annotation is provided in Supplementary Table S3. A comprehensive list of all annotated proteins is provided in Supplementary Table S4.

### 3.3 Enzymatic activity profiling

Investigating the venom composition and bioactivity we adopted and modified previously published protocols for enzymatic activity assays on protease, PLA_2_, FXa-like, thrombin-like, and plasmin-like activities (Schulte *et al*., 2024; Avella *et al*., 2024), as well as effects on mammalian cell viability (Hurka *et al*., 2022) and hemolysis (Avella *et al*., 2024). All analyses were performed in at least triplicate. Unless otherwise stated, measured signals at specific wavelengths were averaged (x̅), subtracted by their respective negative control, and normalized against their respective positive control:

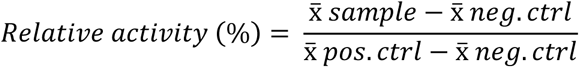

#### 3.3.1 Protease activity assay

The protease activity of the venoms was assessed using the non-specific Protease Activity Assay Kit (Calbiochem, cat. no. 539125) for the 96-well plate format. Venoms were redissolved (Reaction buffer 1:6 ddH_2_O) and added to the substrate (final concentrations 400, 200, 100, 50, and 25 μg/ml), as well as the negative control (Reaction buffer 1:6 ddH_2_O, 0%) and the positive control (166 µg/ml trypsin in Reaction buffer 1:6 ddH_2_O, 100%). After incubation for 2 h at 37 °C and shaking at 120 rpm on a Multitron device (Infors HT), the absorption was measured at λ = 492 nm on a Synergy H4 Hybrid Microplate Reader (BioTek) operated using Gen 5 v2.09 software (BioTek). Raw data from the protease activity assay is provided in Supplementary Table S5.

#### 3.3.2 Phospholipase A_2_ assay

Phospholipase A_2_ activity was assessed using the EnzChek Phospholipase A_2_ Assay Kit (Invitrogen, cat. no. E10217) for the 96-well plate format. Venoms were redissolved in ddH_2_O and mixed with the substrate (final concentrations: 50, 25, 12.5, 6.25, and 3.125 µg/ml), as well as the negative control (Reaction Buffer 1: 20 ddH_2_O, 0%) and the positive control (5 U/ml purified bee venom phospholipase A_2_ in Reaction buffer 1:20 ddH_2_O, 100%). After incubation for 1 h at room temperature, the plates were measured on a Synergy H4 Hybrid Microplate Reader (BioTek) operated using Gen 5 v2.09 software (BioTek), with excitation set to λ = 470 nm and absorbance set to λ = 515 nm. Raw data from the phospholipase A_2_ activity assay is provided in Supplementary Table S6.

#### 3.3.3 Factor Xa-like activity assay

The Factor Xa (FXa)-like activity was assessed using the FXa Activity Fluorometric Assay Kit (MAK238-1KT, Sigma-Aldrich) for the 96-well plate format. Lyophilized venoms were redissolved (Factor Xa Assay Buffer 20: 1 ddH_2_O) and added to the substrate (final concentrations 50, 25, 12.5, 6.25, and 3.125 μg/ml), as well as the negative control (FXa Assay Buffer 1:20 ddH_2_O, 0%) and the positive controls (2 µg/ml FXa Enzyme Standard in FXa Assay Buffer 20:1 ddH_2_O, 100%). The assay was transferred to a Synergy H4 plate reader (H4MLFPTAD, BioTek) operated using Gen 5 v2.09 software (BioTek) for 15 min of incubation at 37 °C, protected from light. The FXa activation was measured, with excitation set to λ = 350 nm and fluorescence detection at λ = 450 nm. Raw data from the FXa activity assay is provided in Supplementary Table S7.

#### 3.3.4 Thrombin-like activity assay

The thrombin-like activity was assessed using the Thrombin Activity Fluorometric Assay Kit (Sigma-Aldrich, MAK242) for the 96-well plate format. Venoms were redissolved (Thrombin Assay Buffer 1:20 ddH_2_O) and added to the substrate (final concentrations 50, 25, 12.5, 6.25, and 3.125 µg/ml), as well as the negative control (Thrombin Assay Buffer 1:20 ddH_2_O, 0%) and the positive control (0.3 µg/ml Thrombin Enzyme Standard in Thrombin Assay Buffer 20:1 ddH_2_O, 100%). In a standardized pre-measurement procedure, the assay was handled for 5 min at room temperature to define a replicable starting time point, before being transferred to a Synergy H4 plate reader (H4MLFPTAD, BioTek) operated using Gen 5 v2.09 software (BioTek). After 15 min of incubation at 37 °C, protected from light, the thrombin activation was measured, with excitation set to λ = 350 nm and fluorescence detection at λ = 450 nm. Raw data from the thrombin activation assay is provided in Supplementary Table S8.

#### 3.3.5 Plasmin-like activity assay

The plasmin-like activity was assessed using the Plasmin Activity Assay Kit (MAK244, Sigma-Aldrich) for the 96-well plate format. The lyophilized venoms were suspended in Plasmin Assay Buffer 20:1 ddH_2_O) and added to the substrate (final concentrations 50, 25, 12.5, 6.25, and 3.125 μg/ml), as well as the negative control (Plasmin Assay Buffer 1:20 ddH_2_O, 0%) and the positive controls (5 µg/ml Plasmin Enzyme Standard in Plasmin Assay Buffer 20:1 ddH_2_O, 100%). The assay was transferred to Synergy H4 plate reader (H4MLFPTAD, BioTek) operated using Gen 5 v2.09 software (BioTek), for 15 min of incubation at 37 °C, protected from light. Plasmin activation was measured with excitation set to λ = 360 nm and fluorescence detection at λ = 450 nm. Raw data from the plasmin activation assay is provided in Supplementary Table S9.

### 3.4 Plasma coagulation assays

The assessment of the venom induced coagulation time (VICT), the prothrombin time (PT) and the activated partial thromboplastin time (aPPT), was performed in a setup with platelet-poor plasma and a microplate reader (Pratt and Monroe, 1992) refined for snake venom-induced coagulation studies (Kerns *et al*., 1999; Banerjee *et al*., 2005; Barnwal *et al*., 2016). Human blood was obtained according to DIN 58905 from a healthy volunteer with no diagnosed circulatory or coagulation disease or medication intake before donation. To obtain platelet-poor plasma (PPP), the blood was added (9:1) to a sterile-filtered sodium citrate solution (0.11 mol/l) and centrifuged for 15 min at 2500 × *g*. The supernatant, essentially platelet-free plasma, was aliquoted and used immediately or flash-frozen in liquid nitrogen to be stored at -80 °C until further use. Preliminary to the assay, PPP aliquots were thawed and kept at room temperature at a maximum of 4 h. Setting up the assay, 25 µl of each PPP and lyophilized venom (4x final concentration of 10, 1, 0.1, and 0.01 µg/ml) reconstituted in Tris-Buffered Saline (TBS) were added to a transparent, flat-bottom 96-well plate. Depending on the assessment, 25 µl of the following were added: (i) TBS for the VICT; (ii) TBS with 5% Dade®Innovin (B4212-50, Siemens Healthcare) for the PT; (iii) DAPTTIN®TC (5035060, Technoclone) for aPPT. Plates were transferred to a microplate reader (CLARIOstar Plus, BMG Labtech) with the thermostat set at 37 °C and absorbance monitored at λ = 405 nm in 5 s intervals for 2 min before 25 µl of CaCl_2_ (25 mM, final concentration: 6.25 mmol/l) were injected automatically via the internal injection system (t = 0) and monitoring continued for 8 min. The controls included a venom-free sample for each assessment, a sample without Dade®Innovin for the VICT and PT, and a sample without CaCl_2_ injection for the aPPT, substituted each by 25 µl TBS. There is no universally accepted standardized method to determine optically measured coagulation time, so we used the time point when maximum fibrin deposition occurs. This is identical to the highest change of absorption signal. This was determined by calculating the first derivative of absorbance at λ = 405 over time:

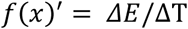

*E* = Absorbance (λ = 405 nm)

*T* = Time (s)

Raw and processed data on venom-induced coagulation-, prothrombin- and activated partial thromboplastin time assay are provided in Supplementary Table S10-S13.

### 3.5 Cytotoxic activity profiling

#### 3.5.1 Cell viability assay

The cytotoxicity of venoms was assessed in Madin-Darby canine kidney II (MDCK II; kindly provided by Prof. Dr. Eva Böttcher-Friebertshäuser, Institute of Virology, Philipps University, Marburg) and human epithelial lung adenocarcinoma (Calu-3; SCC438, Merck) cell lines using the CellTiter-Glo Luminescent Cell Viability Assay (G7570, Promega). The venoms were redissolved in cultivation medium and added to the plate (final concentrations 50, 25, 12.5, 6.25, and 3.125 µg/ml), as well as the negative control (cultivation medium, 100% cell viability) and the positive control (100 µM ionomycin in DMSO, 0% cell viability). Confluent cells were treated with the venom/controls and incubated at 37 °C in a 5% CO_2_ atmosphere. After 48 h, luminescence was measured according to the manufacturer’s protocol on a Synergy H4 Hybrid Microplate Reader operated using Gen 5 v2.09 software (BioTek). Raw data from the cell viability assay is provided in Supplementary Table S14.

#### 3.5.2 Hemolytic activity assay

The hemolytic activity was assessed in Dulbecco’s phosphate-buffered saline (DPBS) and additional DBPS with supplemented Ca^2+^ and Mg^2+^ (DPBS+), the latter to saturate potential cofactors. Lyophilized venom was redissolved in the respective buffer and added to the purified erythrocytes (final concentrations 50, 25, 12.5, 6.25, and 3.125 μg/ml), as well as the negative control (DPBS/DPBS+ 20:1 ddH_2_O, 0% lysis) and the positive control (1% Triton X-100 in DPBS/DPBS+ 20:1 ddH_2_O, 100% lysis). After incubation at 37°C for 1 h, the absorbance was detected at λ = 405 nm with a Synergy H4 plate reader (H4MLFPTAD, BioTek) operated using Gen 5 v2.09 software (BioTek). Raw data and buffer receipt from the hemolytic assay are provided in Supplementary Table S15.

### 3.6 Data visualization

The distribution shown in Figure 1 was generated in RR (R Core Team, 2024) using the following packages: tmap (Tennekes, 2018), sf (Pebesma, 2018; Pebesma and Bivand, 2023), dplyr (Wickham *et al*., 2023), magrittr (Bache *et al*., 2022), purr (Wickham *et al*., 2025), raster (Hijmans *et al*., 2025), rnaturalearth (Massicotte *et al*., 2023), rnaturalearthdata (South *et al*., 2024), rnaturalearthhires (South *et al*., 2025), and elevatr (Hollister *et al*., 2023). The chromatographic profiles, proteomic data, and bioactivity results were visualized using Origin 2020b.

## 4 Results

### 4.1 Venom yield

The sampled *V. berus* specimens measured from 50 to 59 cm in total length, and their venom yields ranged between 0.783 and 9.708 mg (dry weight) (Figure 2). Females were generally larger than males (average: male 53 cm vs. female 57 cm) and usually provided higher amounts of venom (average: male 2.380 mg vs. female 4.436 mg). In general, the amount of retrieved dry venom tended to increase with the specimen’s body size.

**Figure 2.**
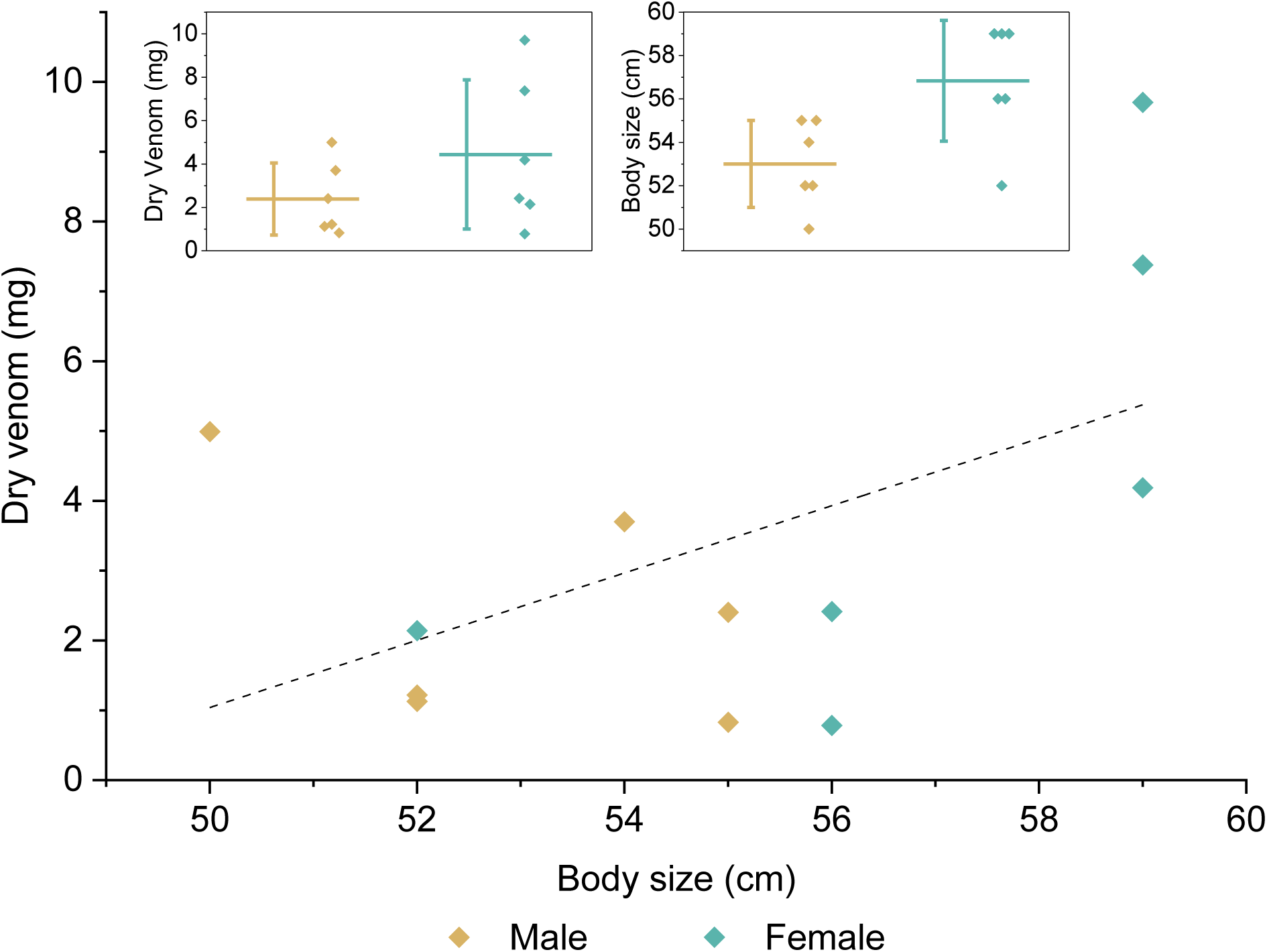
Body size positively correlates with venom yield in *Vipera berus*. The graph shows the correlation of the specimen’s size and amount of dry venom for male and female *V. berus*. The linear fitting curve is represented as a dashed line. The insets display the distribution, mean value, and standard distribution of measured dry venom (upper) and body size (lower) separated by sex.

### 4.2 Compositional profiling of *Vipera berus* venoms

To investigate compositional differences between the *V. berus* venoms from specimens of different sexes, we analyzed their electrophoretic profiles obtained by performing SDS-PAGE under reducing and non-reducing (Figure 3 A, B) conditions. We assigned the obtained bands to putative toxin families based on the information reported in previous works (Latinović *et al*., 2016; Bocian *et al*., 2016; Al-Shekhadat *et al*., 2019).

**Figure 3.**
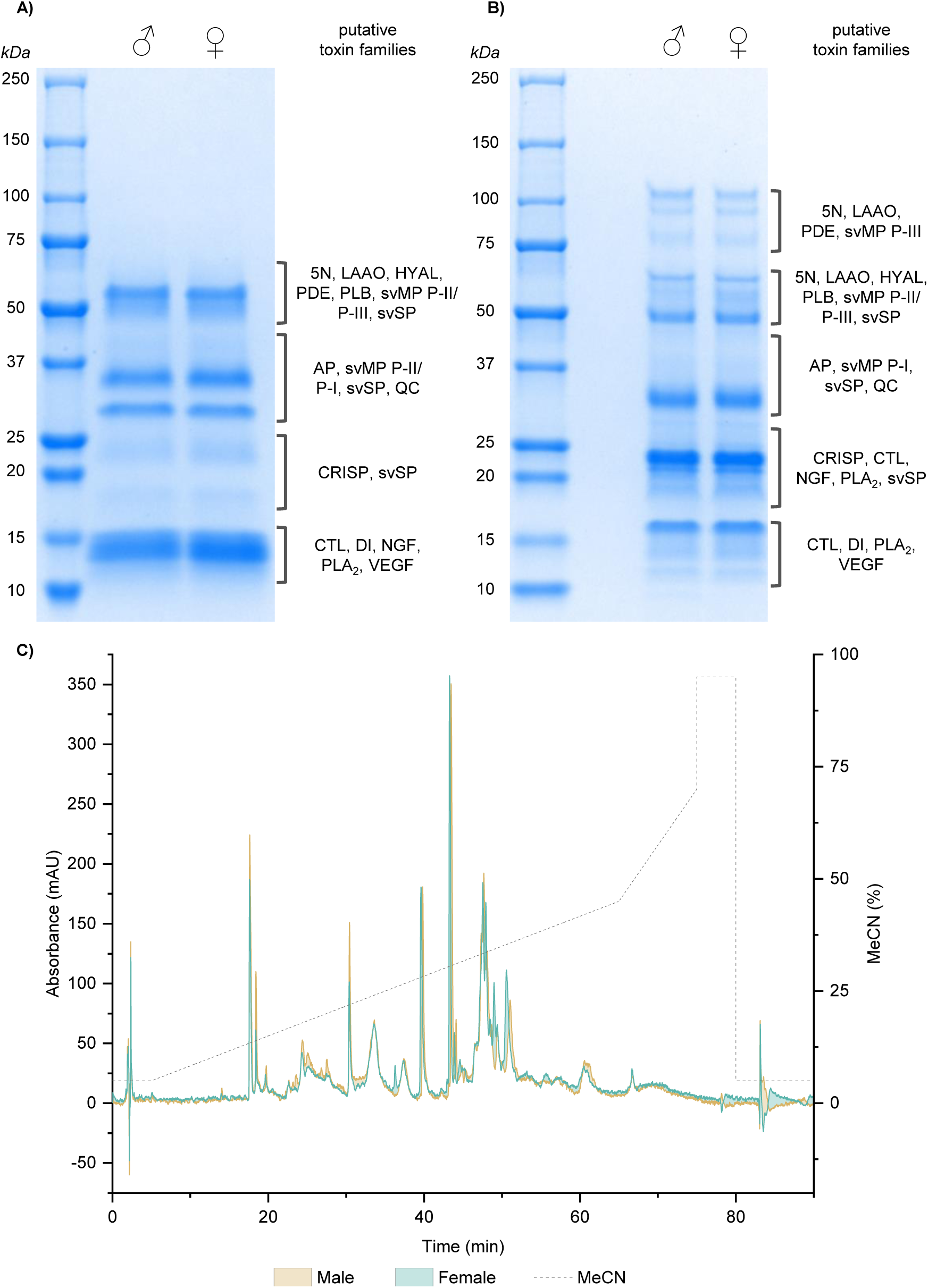
Venom profiling of adult male and female *Vipera berus* specimens. SDS-PAGE profiling under A) reduced and B) non-reduced conditions, and C) RP-HPLC profile (absorbance (mAU) at λ = 214 nm)). The initial peak (0) corresponds to the sample injection. Abbreviations: 5N, 5′-nucleotidase; AP, aminopeptidase; CTL, C-type lectin, snaclec, and C-type lectin-related protein; HYAL, hyaluronidase; LAAO, L-amino acid oxidase; NGF, nerve-growth factor; PLA2, phospholipase A2; PLB, phospholipase B; PDE, phosphodiesterase; QC, glutaminyl cyclase; svMP, snake venom metalloproteinase; svSP, snake venom serine protease; VEGF, vascular endothelial growth factor.

The SDS-PAGE at reducing conditions (Figure 3 A) revealed that for both venom pools the protein mass ranges from 12 to 70 kDa. The most intense bands cluster at 13-15 kDa, 30 kDa, 35 kDa, and 55-60 kDa. Less conspicuous bands are detectable at 12 kDa, 18 kDa, 23-25 kDa, 35-43 kDa, 48-55 kDa, and 60-70 kDa. While differences in band diversity are not apparent, the bands at 13 kDa and 23 kDa appear slightly more intense in the female sample. Under non-reducing conditions (Figure 3 B), the protein masses range from 12 to 115 kDa for both venoms. The most intense bands appear to be at 16 kDa, 23 kDa, 24 kDa, 32 kDa, and 50 kDa. Less intense bands are visible at 12 kDa, 13-15 kDa, 19-22 kDa, 26-29 kDa, 55-60 kDa, 60-65 kDa, 80-85 kDa, 90-95 kDa, and 110-120 kDa. No dominant differences in band diversity are evident, but the bands at 25 kDa and 60 kDa appear slightly more intense in the female venom.

The chromatograms of the venoms present 45 peaks in total. Both profiles appear largely similar, and only minor differences could be detected. For instance, peak 20 is seemingly absent in the male venom. Additionally, peaks 2, 3, 7, 8, 9, 10, 11, 12, 14, 15, and 16 are slightly higher in the male venom, whereas peaks 33, 34, 35, 36, 37, 41, and 44 are slightly higher in female venom.

Furthermore, we assessed for qualitative differences in venom composition via shotgun proteomics using DDA and DIA mass spectrometry (Figure 4). In DDA, we annotated a total of 179 proteins, of which 106 were shared between both sexes. 36 were exclusively found in male venom and 37 in female venom. (Figure 4, C) Those proteins were further clustered into protein groups and annotated. 97 protein groups were identified in the male venom (Figure 4, A), and 99 in the female venom (Figure 4, B). In DIA mode, 265 total proteins were annotated, with 153 shared between males and females. 60 were only found in male venom and 52 only found in female venom (Figure 4, C). Clustering into protein groups resulted in 141 hits for male venom (Figure 4, D) and 134 for female venom (Figure 4, E).

**Figure 4.**
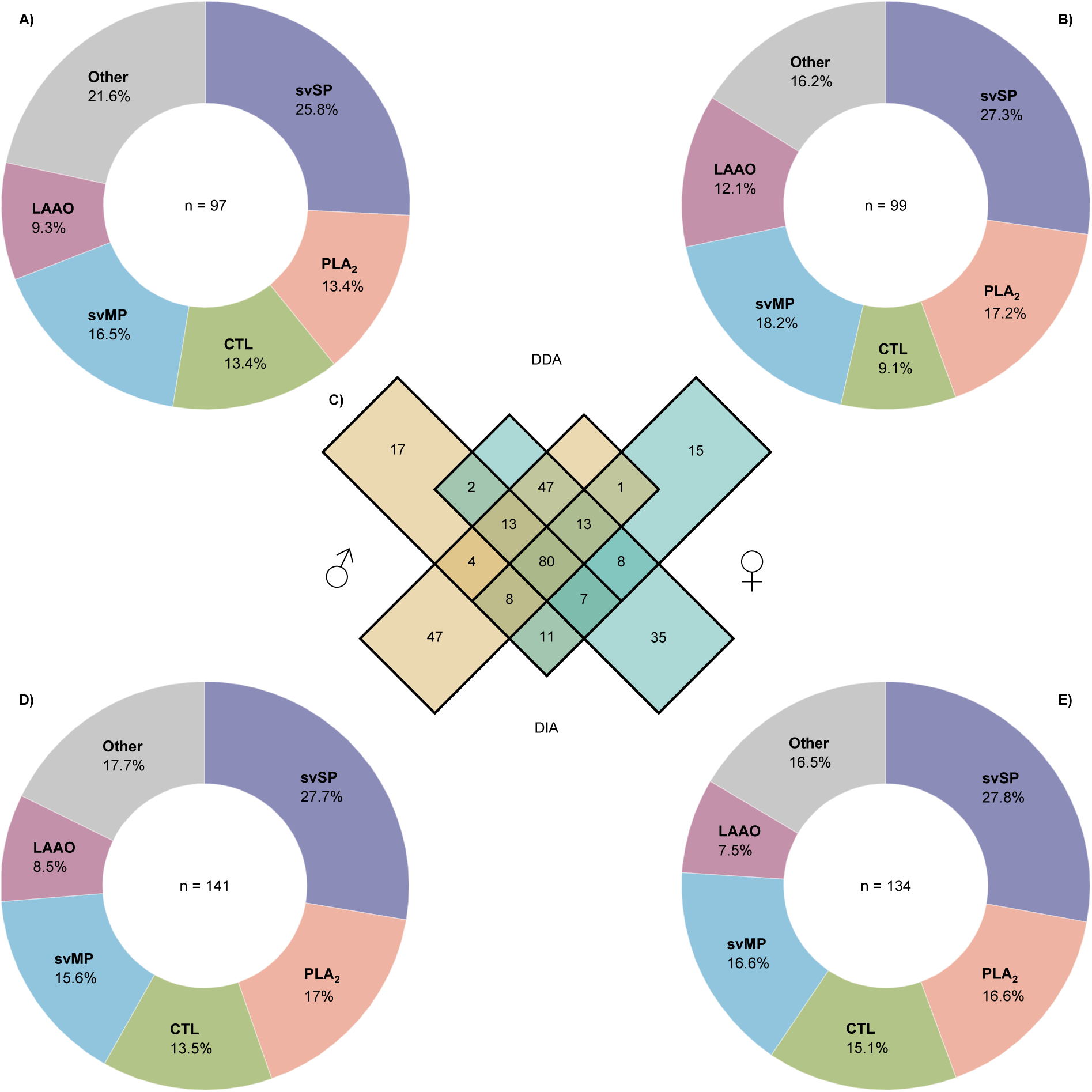
Comparative shotgun proteomics of the venom of adult male (A & D) and female (B & E) *Vipera berus.* Proteomic data was retrieved in data-dependent acquisition (DDA, A & B) and data-independent acquisition (DIA, D & E) mass spectrometry mode, respectively. The Venn diagram C) displays the total annotated proteins, exclusively for each approach and shared between approaches. Furthermore, qualitative venom proteomes (B-E) are presented with the relative diversity of protein families for the respective approach, based on the total number of identified protein groups (n). Abbreviations: svSP, snake venom serine protease; PLA2, phospholipase A2; svMP, snake venom metalloproteinase; CTL, C-type lectin inclusive C-type lectin-like protein and snaclec; LAAO, L-amino acid oxidase. “Other” includes protein families that comprise less than 5% of total protein family diversity: CRISP, cysteine-rich secretory proteins; KUN, Kunitz-type inhibitor; DI, disintegrin; PDE, phosphodiesterase; AP, aminopeptidase; PLB, phospholipase B; VEGF, vascular endothelial growth factor F; 5N, 5’-nucleotidase; NGF, nerve growth factor; HYAL, hyaluronidase; QC, glutaminyl cyclase; C3, venom complement C3 homolog; NP, natriuretic peptide; UNSPEC, venom non-specific component.

Subsequent classification of protein groups revealed that five protein families, svSPs, PLA_2_s, svMPs, C-Type lectins (CTLs, including C-type lectin related proteins and snaclecs), and L-amino acid oxidases (LAAOs), contain more than 5% of total protein family diversity in both sexes. In all cases, svSPs show the highest diversity of venom components (Male: DDA 25.8%, DIA 27.7%; Female: DDA 27.3%, DIA, 27.8%). In DDA, the non-enzymatic CTLs follow next in diversity for both sexes (male 16.5% and female 18.2%), continued in male venom by PLA_2_s and svMPs equally (13.4%), as well as LAAOs (9.3%), while in female venom PLA_2_s (17.2%) and LAAOs (12.1%) precede svMPs (9.1%) in diversity. In DIA, regardless of the venom, svSPs are followed by PLA_2_s (male 17.0% and female 16.6%), svMPs (male 15.6% and female 16.6%), CTLs (male 13.5% and female 15.1%), and LAAOs (male 8.5% and female 8.5%). The diversity dominating protein families make up 78.4% for DDA and 82.3% for DIA mode in male venom, and 83.8% for DDA and 83.5% for DIA in female venom. Protein families with generally less than 5% of the total diversity share (“Others”) made collectively the greatest variation between the sexes in DDA mode. In general, differences between sexes in protein family diversity showed a lesser extent in DIA mode compared to DDA mode, comparing the proportions of major protein families.

Overall, our SDS-PAGE and RP-HPLC profiling as well as our proteomic analysis under different data acquisition modes, revealed that the venom profiles of male and female *V. berus* are almost identical. Small deviations occur but only manifest in minuscule differences or affect minor components.

### 4.3 Bioactivity profiling

To investigate functional differences between male and female venoms, we compared their normalized enzymatic activity and effects on mammalian cell lines and erythrocytes (Figure 5). Venoms were tested at concentrations of 50, 25, 12.5, 6.25, and 3.125 µg/ml in every assay, except for the protease activity assay, where concentrations of 400, 200, 100, 50, and 25 µg/ml were applied.

**Figure 5.**
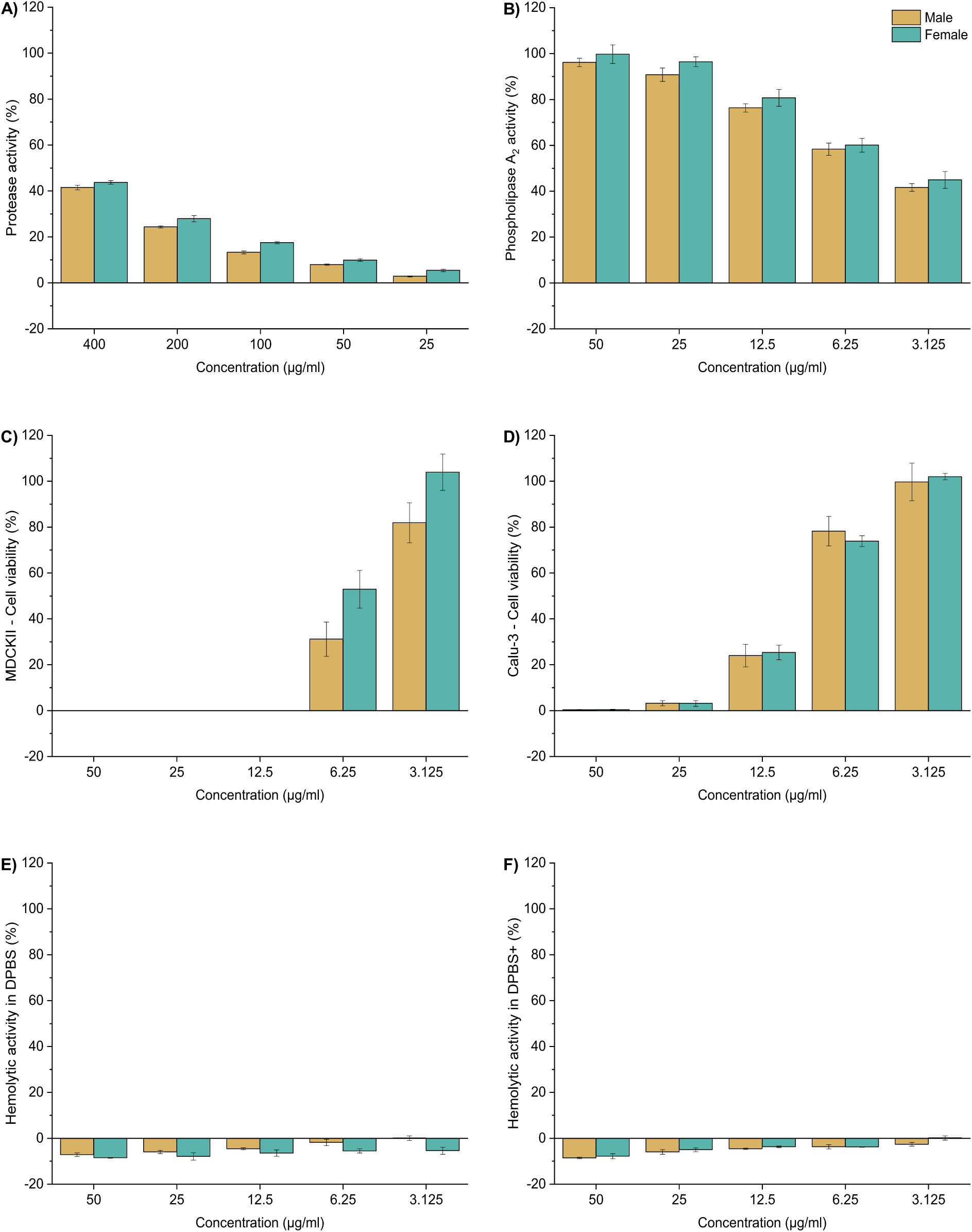
Comparison of venom bioactivities of adult male and female *Vipera berus* venoms. A) Protease activity; B) Phospholipase A2 activity; venom-induced effect on the cell viability of C) Madin-Darby canine kidney II (MDCK II) cell line, and D) human epithelial lung adenocarcinoma (Calu-3) cell line; venom-induced hemolysis on equine erythrocytes in E) Dulbecco’s phosphate-buffered saline (DPBS) and F) DBPS with supplemented Ca2+ and Mg2+ (DPBS+). Displayed are the results at five concentrations. Data are means ± standard deviations of technical triplicates (n = 3).

Protease activity of both venoms (Figure 5, A) showed a concentration-dependent effect, ranging between 44% and 3%. Measured activity between male and female venoms was highly similar, e. g. 44% (female) vs. 42% (male) at 400 µg/ml, or 28% (female) vs. 24% (male) at 200 µg/ml. Likewise, the measured PLA_2_ activity (Figure 5, B) showed a concentration-dependent effect for both samples, ranging from 100% to 42%. Again, measured differences were only marginal at every tested concentration, e.g.: 100% vs. 96 at 50 µg/ml, or 96% vs. 91% at 25 µg/ml in female and males, respectively.

Reminiscent of the enzymatic activity spectrum, effects from male and female venoms on the viability of MDCK II cells (Figure 5, C) were comparable at all tested concentrations. No cytotoxicity was recorded at lower concentrations. The effects on Calu-3 viability were similar at all concentrations (Figure 5, D). Both venoms were strongly cytotoxic to a concentration of 12.5 µg/ml (24-25% cell viability) with marginal cytotoxicity at 6.25 µg/ml (78-74% cell viability) and no effect was detected at lower concentrations. Relevant hemolytic activity could not be measured up to the maximal concentration of 50 µg/ml for neither male nor female venom against purified equine erythrocytes (Figure 5 E, F).

Furthermore, we compared the effects of our venom samples regarding their ability to target the coagulation cascade (Figure 6, A-C). We tested FXa-like-, thrombin-like-, and plasmin-like activities, as well as the venoms’ general coagulation capability by venom-induced coagulation time, prothrombin time, and activated partial thromboplastin time. No relevant FXa-like activity (Figure 6, A) was observed for any of the two venom pools, regardless of the tested concentrations. The thrombin-like activity (Figure 6, B) of both venoms displayed a similar, potent concentration-dependent effect, ranging from 138% (Female 50 µg/ml) to 33% (Male 3.125 µg/ml), exceeding the positive control ≥ 12.5 µg/ml. Similarly, plasmin-like activity (Figure 6, C) also showed a highly potent, concentration-dependent effect. Again, both venoms exhibited similar activities with minor differences, exceeding the positive control at concentrations ≥ 6.25 µg/ml.

**Figure 6.**
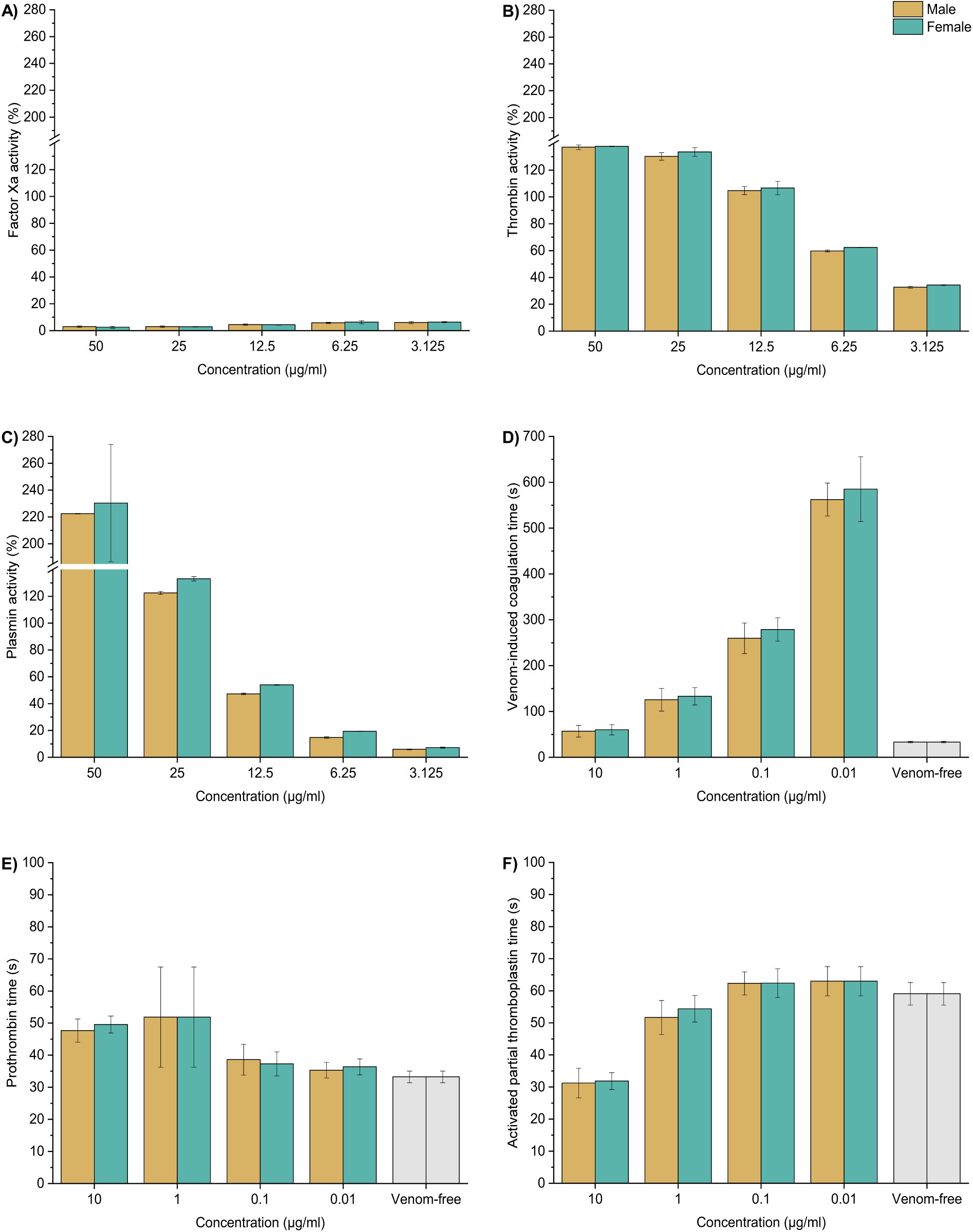
Effects on coagulation exerted by adult male and female *Vipera berus* venom: Displayed is the enzymatic activity in the coagulation cascade with A) Factor Xa (FXa)-like activity, B) Thrombin-like activity, and C) Plasmin-like activity. Further is presented the impact on coagulation times in human platelet-poor plasma with the D) Venom-induced coagulation time; E) Prothrombin time; F) Activated partial thromboplastin time. The maximum coagulation at each concentration, compared to the respective positive control, is defined as the highest change in fibrin deposition over time, measured as absorbance at λ = 405. Data are means ± standard deviations of technical triplicates (n = 3, A-C) and octuplicates (n = 8, D-F).

In addition, we assessed for differences in the coagulation time of human PPP for VICT, PT, and aPPT (Figure 6, D-F). The VICT assay (Figure 6, D) displays similar coagulation times, occurring in a dose-dependent manner (10 µg/ml: male 57 s, female 60 s; 1 µg/ml: male 126 s, female 133 s; 0.1 µg/ml: male 260 s, female 279 s; 0.01 µg/ml: male 562 s, female 585 s). Assessing PT (Figure 6, E), a reference for coagulation-induction through the extrinsic coagulation pathway, neither male and nor female venom had a strong effect on the coagulation time compared to the control (10 µg/ml: male 48 s, female 50 s; 1 µg/ml: male 52 s, female 52 s; 0.1 µg/ml: male 39 s, female 37 s; 0.01 µg/ml: male 35 s, female 36 s). In the aPPT assay (Figure 6, F), assessing for coagulation induction via the intrinsic pathway, both venoms again show similar coagulation times with stagnating dose-dependency and very similar effects (10 µg/ml: male 31 s, female 32 s; 1 µg/ml: male 52 s, female 54 s; 0.1 µg/ml: male 62 s, female 62 s; 0.01 µg/ml: male 63 s, female 63 s). At the highest concentration evaluated, a strong decrease in aPPT could be observed. However, no variation between sexes was determined.

## 5 Discussion

### 5.1 Low extent of sex-based venom variation in *Vipera berus*

As other functional traits relevant to an animal’s fitness, snake venoms are shaped by selective pressures and environmental cues (Casewell *et al*., 2013; Schendel *et al*., 2019). Venoms generally serve the three primary functions of intraspecific interaction, defense, and predation (Schendel *et al*., 2019). However, the trophic role of venom appears to be the most critical function in snakes, while defense appears to be pivotal only in few lineages (Ward-Smith *et al*., 2020; Kazandjian *et al*., 2021) and intraspecific interaction has not yet been shown to play a role (Daltry *et al*., 1996b; Casewell *et al*., 2013). That said, identifying the factors determining the occurrence of intraspecific venom variation in snakes and contextualizing them within an ecological framework is crucial for understanding venom evolution as well as improving snakebite treatment. Particularly, intersexual size differences have been proposed to drive venom variation in several taxa (e.g., *Calloselasma rhodostoma* (Daltry *et al*., 1996a), *B. jararaca* (Menezes *et al*., 2006; Zelanis *et al*., 2016), *B. moojeni* (Hatakeyama *et al*., 2021; Ferreira-Rodrigues *et al*., 2024), and *B. leucurus* (Machado Braga *et al*., 2020)). Differences in body size ultimately cause differences in gape size, and hence influence the ability to swallow prey (Arnold, 1993). Such discrepancies between male and female conspecifics are often associated with dietary preferences between sexes (Nogueira *et al*., 2003; Sasa *et al*., 2009). Hence, intersexual venom variation has typically been explained with the need to target different prey spectra as a result from the physical constraints stemming from size differences (Menezes *et al*., 2006; Zelanis *et al*., 2016). However, to date a relatively limited number of studies has explored the frequency and extent of sex-related differences in snake venoms. *Vipera berus*, a snake that has been shown to exhibit sexually dimorphism and to display venom variation, has seldomly been studied for the role of sex in manifesting aberrant venom profiles.

We determined the venom profiles of male and female *V. berus* from a German population. The SDS-PAGE showed no differences in band patterning between sexes, although slightly increased intensity of certain bands could be detected in female venoms. Nonetheless, considering earlier reported discrepancies in SDS-PAGE venom profiles between individual Hungarian *V. berus* specimens (Malina *et al*., 2017), this variation seems unlikely to be strictly sex-based and likely falls within the individual venom variation already detected in this species. Similarly, our HPLC profiles present highly similar peak landscapes for both sexes, suggesting qualitatively and quantitatively similar venom profiles. Furthermore, we performed shotgun proteomics, revealing svSPs, PLA_2_s, CTLs, svMPs, and LAAOs as major diverse components of venoms from the analyzed specimens. We also detected considerable overlap of hits retrieved from publicly available databases between both samples, suggesting minor variability between the male and the female venoms tested. These proteomic findings are in agreement with the low extent of venom variation already seen by our SDS-PAGE and RP-HPLC profiles.

We also gathered additional evidence for supporting the similarity of male and female venoms in our functional screening. The results of PLA_2_ and protease activity assays, as well as those regarding the assessment of thrombin-, plasmin- and FXa-like activities, were generally comparable between sexes. The effects on coagulation times are comparable between male and female venoms. Likewise, no relevant differences were detected on mammalian cells. These findings are in line with the results of Schulte *et al*. (2024), comparing the cytotoxic effects of the venoms *V. berus* specimens exhibiting different color phenotypes.

Our data suggests that venom composition and activity seem to be nearly identical between male and female venoms in *V. berus*. Especially when facing the variability of individual venoms (Malina *et al*., 2017), the extent of venom variation observed across our proteomic and functional comparison appears negligible. While we retrieved some smaller differences for some activities, we believe that these differences have little to no role in an ecological or clinical framework since they usually occurred only at the lowest concentrations. We come to this conclusion based on the >10 mg dry venom that was gathered from the analyzed snakes in our study and by facing that *V. berus* can inject up to 10-18 mg (Al-Shekhadat *et al*., 2019). Therefore, it can be expected that the injected amounts in real-world encounters symptomatically overshadow such subtle differences at low dosages. Our data agrees with natural history data, as no intersexual differences in foraging behavior and prey choice are reported for *V. berus* to this day (Forsman, 1991). Therefore, it is likely that both sexes of *V. berus* produce chemically and functionally highly comparable venoms in response to similar prey spectra targeted despite their sexually dimorphic nature.

### 5.2 Clinical considerations and the effect of sex on envenoming

Parts of our work were set out to unveil intersexual differences in venom function and activity of *V. berus* venom. While these revealed no striking differences between the venoms of male and female common adders, this investigation helps to shed new light on the pharmacological effects caused by this species’ venom.

For instance, our work revealed that *V. berus* venom acts very potently on the coagulation cascade and targets several of its elements. Considering the assessment of FXa-like, thrombin-like, and plasmin-like activity, both venoms showed similarly strong effects (Figures 6, B and C). Considering that endogenic thrombin and plasmin are serine proteases, these findings possibly align with the high protease activity measured for *V. berus* venom (Figure 5, A). On the other hand, no FXa-like activity was detectable in either venom at the tested concentrations. The performed plasma coagulation time assays provided further evidence for the ability of *V. berus* venom to disrupt the normal coagulation process of human plasma. The venom was able to induce coagulation in recalcified human plasma in a concentration-dependent manner, with male and female venom possessing comparable potency. At venom concentrations of 10 µg/ml and 1 µg/ml, both male and female venom induced an increase in the prothrombin time (PT) while the aPPT was strongly reduced at the highest concentration tested (10 µg/ml). The increase in PT might be due to the disruption of membranes by Phospholipases in the venom, as the tissue factor used for the PT assay is enveloped in phospholipid vesicles. Considering the observed thrombin-like activity, it has been shown that some svSPs exhibit thrombin-like activity, often recalled as snake venom Thrombin-like enzymes (svTLEs), summarized in fibrino(geno)lytic activity. Such svTLEs have been identified in the proteomic data of both sexes (see Supplementary Table S4). The svTLEs vary in their mode of action; however, in general, they only partially mimic thrombin activity and do not activate further coagulation factors, resulting in incomplete or unstable formation of fibrin clots. Furthermore, some thrombin-like svSPs and svMPs have been shown to act fibrinolytic. This results in a plasmin-like activity as shown in our assessments, degrading the (unstable) fibrin clots (Castro *et al*., 2004; Lu *et al*., 2005; Sajevic *et al*., 2011). This presents the venom as an overall pro-coagulant, acting on the common pathway of the coagulation cascade, by promoting an unstable clot formation and the consumption of coagulation factors. An impaired coagulation at the bite site might cause the described local bleeding due to *V. berus* envenomation (Warrell, 2005; Paolino *et al*., 2020). However, systemic coagulopathy is rarely reported in clinical cases (Persson, 2014; Jollivet *et al*., 2015; Dyląg-Trojanowska *et al*., 2018; Hermansen *et al*., 2019), likely due to an insufficient venom dosage to induce a systemic effect in humans.

Besides the effects on the coagulation cascade, our investigation of cell viability allows us to draw conclusions upon the cytotoxicity of *V. berus* venom. Our experiments showed that male and female venoms had similar effects on mammalian cells. Cell viability of MDCK II cells was not detectable at venom concentration ≥ 12.5 µg/ml, whereas viability of Calu-3 cells was barely detectable at 25 µg/ml venom. In line with our results, *V. berus* venom is known to cause local tissue damage, hemostatic imbalance, and organ damage (Al-Shekhadat *et al*., 2019; Siigur and Siigur, 2022; Shchypanskyi *et al*., 2024). The most common symptoms caused are local swelling and tissue damage and in more severe cases renal damage, which aligns with our results from cell viability assays (Warrell, 2005; Persson, 2014; Valenta *et al*., 2014; Dyląg-Trojanowska *et al*., 2018; Hermansen *et al*., 2019; Paolino *et al*., 2020). The high potency in our cell assays may, at least partially, also explain frequent reports of long-lasting local swelling (Warrell, 2005; Persson, 2014). Concerning the hemolytic activity assay, in line with previous reports (Schulte *et al*., 2024), we did not observe any activity at the tested concentrations. We therefore hypothesize that described symptoms in clinical reports indicating erythrocytopenia (Persson, 2014; Hermansen *et al*., 2019) are secondary effects of *V. berus* venom, e.g. due to hemorrhage due to local tissue destruction and not stemming from direct hemolytic activity.

Our analysis of *V. berus* venom demonstrates strong effects on the coagulation cascade. It causes disturbances in the common coagulation pathway, likely caused by its thrombin-like and plasmin-like activity. It also has noteworthy effects on cell viability, providing the mechanistic basis for the clinically observed tissue damage. The major enzymatic activities are protease and PLA_2_ activity, both of which are considerable in this species. Interestingly, while the venom profiles and activities between males and females were found to be highly similar, there might be a clinically relevant effect of sex. In virtually all conducted assays, we retrieved very clear dose-dependencies. When analyzing our venom yields, gathered by mimicking defensive bites, we found that females were capable of delivering much higher amounts of venom. Ergo, while venom biochemistry is conserved between both sexes, female *V. berus* may bear a higher potential in causing severe envenoming by delivering bites with a higher dosage of venom.

### 5.3 Novel proteomic perspectives on *Vipera berus* venom

*Vipera berus* is Earth’s most widespread medically relevant snake, and is responsible for the highest fraction of snakebites in Europe (Paolino *et al*., 2020). Surprisingly little is known about its venom composition and activity from across its range. Our investigation of sex-based venom variation allows us to gather important new insights into the venom of this severely overlooked snake.

Most importantly, our work provides the first venom proteomes for *V. berus* specimens from Germany, and Central Europe in general. Previous studies on the venom of *V. berus* were mostly conducted on specimens from Eastern and Southern Europe. They identified a similar set of major components, such as svSPs, PLA_2_s, and svMPs. The families CTL and LAAO occur consistently but in varying proportions in the total venom proteomes (Latinović *et al*., 2016; Bocian *et al*., 2016; Al-Shekhadat *et al*., 2019; Damm *et al*., 2024). A recently published quantitative venom proteome of Norwegian *V. berus* (Nicolaysen *et al*., 2024) presented LAAOs as the most abundant venom components, followed by svMP and svSP. Overall, *V. berus* venom appears to comprise the same major building blocks across the specieś range, all belonging to the classical major viperine venom components *sensu* Damm et al. (2021). However, the relative contribution of each compound appears to differ between localities, probably in response to locally varying selective pressures, or based on different methods. Nonetheless, many populations of Western and Central European countries, as well as of Central and East Asia have never been investigated, and the whole spectrum of *V. berus* venom components and properties is yet to be fully understood.

## 6 Conclusion

Animal venom is a highly functional ecological trait, complex and dynamic between and within taxa. Especially snakes have been shown to produce intraspecifically diverse venom profiles, affecting the mode of action and symptoms of envenomation. Therefore, it is of great interest to unravel the factors shaping venom composition and bioactivity in the context of the evolution of the snake’s ecological niche and to improve the treatment of snake bite patients. Among others, the snake’s sex, in the context of sexual dimorphism, has been discussed. It emerged to be an important factor influencing venom composition in some snakes but overall received comparatively little scientific attention. Using *V. berus*, a promising model snake to study venom variation, our study provides strong evidence for a low extent of sex-based venom variation in this species. While shedding new light on its functional and compositional phenotype via proteomics, profiling, and bioassays, we did not retrieve striking differences between male and female venoms for having an ecologically or medically relevant impact. However, since our work is limited by low sample sizes and only considers a single population, further comparative studies on a larger scale and covering multiple localities are desirable. As obtaining relevant sample sizes is often a considerable hurdle, especially when working with rare, heavily protected, or dangerous species, we acknowledge that this metric cannot always be optimized to the desired extent, such as in the present study. Furthermore, it is worth noticing that the term ‘venom variation’ has so far been applied quite subjectively and in a non-standardized way to address both subtle variations in SDS-PAGE band patterns to dramatic differences in the relative abundances of major venom protein families. Additionally, while some works used rigorous methodological approaches, statistics, and large sample sizes, others were based on less stringent frameworks. As our study highlights, different approaches generating qualitative venom proteomes can have a marked impact on the extent of observed differences in protein diversity within the same sample. We believe that the field of snake venom research would tremendously benefit from a unified definition of best practices and methodological standards to address snake venom variation.

## 7 Supplementary material

Table S1 – Metrics

Table S2 - RP-HPLC

Table S3 - Proteomics settings

Table S4 - Proteomics data

Table S5 - Protease activity

Table S6 - PLA2 activity

Table S7 - Factor Xa activity

Table S8 - Thrombin activity

Table S9 - Plasmin activity

Table S10 - Coagulation summary

Table S11 - VICT

Table S12 – PT

Table S13 - aPPT

Table S14 - Cell viability

Table S15 - Hemolytic activity

## 9 Acknowledgements

We thank the members of the “Terrarienclub Bayreuth und Umgebung e.V.” for donating the venom samples used in this study. We highly appreciate the intellectual support given by various members of the Venture for Interconnection, Protection, Education and Research in Adders, VIPERA e.V.. A.V. acknowledges generous funding from the Hesse Ministry of Science and Art (HMWK) via the LOEWE Centre for Translational Biodiversity genomics. IA, MD, TL, and LS are funded by the Deutsche Forschungsgemeinschaft (DFG, German Research Foundation; refs. 545040837 (IA), 540833593 (MD), and 505696476 (TL and LS), respectively). JE and KH are supported by the BMBF project ASCRIBE (Grant number 01KI2024).

